# Fully Automated and Scalable Pipeline for Macaque Brain Registration

**DOI:** 10.1101/2025.07.10.663842

**Authors:** Junjie Zhuo, Zhanbo Zhang, Chi Xiao, Mingli Wang, Shukang Bi, Liang Shi, Qingming Luo, Hui Gong, Zhiming Shen, Xiaoquan Yang, Pengcheng Li

## Abstract

With the rapid development of various whole-brain ex vivo imaging technologies, there is a growing need to develop cross-modal 3D image registration methods to integrate multimodal imaging datasets. While comprehensive cross-modality registration tools such as D-LMBmap and mBrainAligner have been successfully implemented for mouse brains, macaque whole-brain registration presents unique challenges. These include more pronounced non-uniform deformations in larger ex vivo specimens, greater modality-specific contrast differences relative to standard space, and increased inter-individual variability. To address these challenges, we developed Macaca-Star, which incorporates deep learning models and self-individual MRI to tackle cross-modal and ex vivo sample deformation challenges in macaque whole-brain registration. Macaca-Star provides fully automated alignment of fMOST and 2D fluorescent slice images to the NMT MRI standard space, allowing for comprehensive integration of anterograde axonal projections and retrograde-traced neuronal soma profiles.

## Introduction

The brain, as the most complex organ, requires us to understand it at different scales using various technological methods^1^. Due to the rapid advancement of various ex vivo imaging techniques and the implementation of national brain initiatives^2–4^, we are currently generating vast amounts of neuroimaging data that reflect information such as cytoarchitecture^5, 6^, connectivity^7, 8^, and spatial transcriptional^9, 10^ information. As more data is generated, an important goal of brain research is to integrate information on cytoarchitecture, connectivity, and gene expression information at various scales and levels to enhance our understanding of brain mechanisms^1, 11^. The key to this integration lies in cross-scale and cross-modal image registration^12, 13^, which unifies the data into a common reference space^14^.

In response to this demand, with the widespread application of deep learning techniques for biomedical imaging analysis, several automated methods have been successfully developed for mice. For instance, Deepslice^15^ allows rapid registration from 2D slices to the 3D CCF standard space^14^. DeepMapi^16^, mBrainAligner^13^, and D-LMBmap^12^, through different deep learning combined with annotation-based strategies, have achieved whole-brain level automatic registration, aligning neuronal projection information, cell distribution, and soma location to the CCF standard space in mice. However, achieving fully automated registration of larger-volume non-human primate and human optical brain sections into common space has remained a challenge. In marmoset, researchers have combined ex vivo and in vivo MRI to achieve registration of histological slices to the standard space^17^. Julish-brain^6^ has achieved registration of 20μm thickness human brain optical slices^18^ to the standard space, but manual intervention is required for correction.

Registration of ex vivo whole-brain optical slice images to the standard space presents three challenges in monkey compared to mouse. First, scale differences. The volume of a mouse brain is nearly 200 times smaller than that of a monkey brain^19^. As the sample volume increases, the deformation differences caused by ex vivo brain tissue processing also become more significant. Second, modality differences. The CCF template in mice^14^ was constructed based on autofluorescence images, whereas there is no standard space generated from optical images for macaque brains. The NMT standard space for macaque^20^ was constructed based on MRI, leading to significant differences in image features such as resolution and texture between modalities. Third, individual differences. It is well-known that individual morphological differences in monkey brains are much greater than those in mouse brains.

In this study, we develop Macaca-Star (Style transfer-based automated registration pipeline for Macaque), which incorporate deep learning model and self-individual MRI to address the cross-modal and ex vivo sample deformation challenges in macaque whole-brain registration. The core idea involves two main points: first, using a style transfer deep learning network^21^ to transform preprocessed 3D whole-brain anatomical feature optical images, such as fMOST propidium iodide (PI) or Blockface image, into MRI image style to reduce modality differences. Second, during the registration procedure, we incorporated self-derived MRI images to maintain the original in vivo structural and morphological characteristics of the macaque brain, thereby minimizing the influence of postmortem tissue deformation and inter-individual variability. With the integration of these two points and the use of symmetric diffeomorphic normalization (Syn) registration methods^22^, we have achieved a fully automated registration pipeline. We have uploaded the relevant code on website (http://atlas.brainsmatics.org/a/zhuo2504).

## Result

### A complete pipeline for whole-brain analysis

Macaca-Star is a fully automated pipeline that can register high-resolution macaque brain slices (∼3 μm thickness in the z axis), such as fMOST, and classical section slice images (40∼240 μm thickness in the z axis) to the NMT standard space. Through the fMOST PI staining channel, we can map the cytoarchitecture information of the PI images, as well as neuronal projection information from other GFP channels, to the NMT standard space. Additionally, Block face images can be used as a bridge to map the cytoarchitecture information or tracing soma locations from 2D stained slices to the 3D standard space. After undergoing independent preprocessing pipelines, different types of images can each undergo the subsequent registration process. We will demonstrate the entire registration process from 2D optical slices to the 3D MRI standard space using typical fMOST anterograde axonal projections and thick-section retrograde tracing neuronal soma images as examples.

Due to the differences in image features (such as brightness, texture, resolution, etc.) and anatomy (including ex vivo slice distortions and individual variations) between macaque optical slices and MRI templates, we designed three modules. These modules are: First, a preprocessing module that reconstruct 2D slice images into smooth 3D images (Fig. 1a); Second, deep learning style transfer module that transforms 3D optical images into MRI-type images (Fig. 1.b); Third, a self-individual MRI guided registration module (Fig. 1c).

**Fig. 1.**
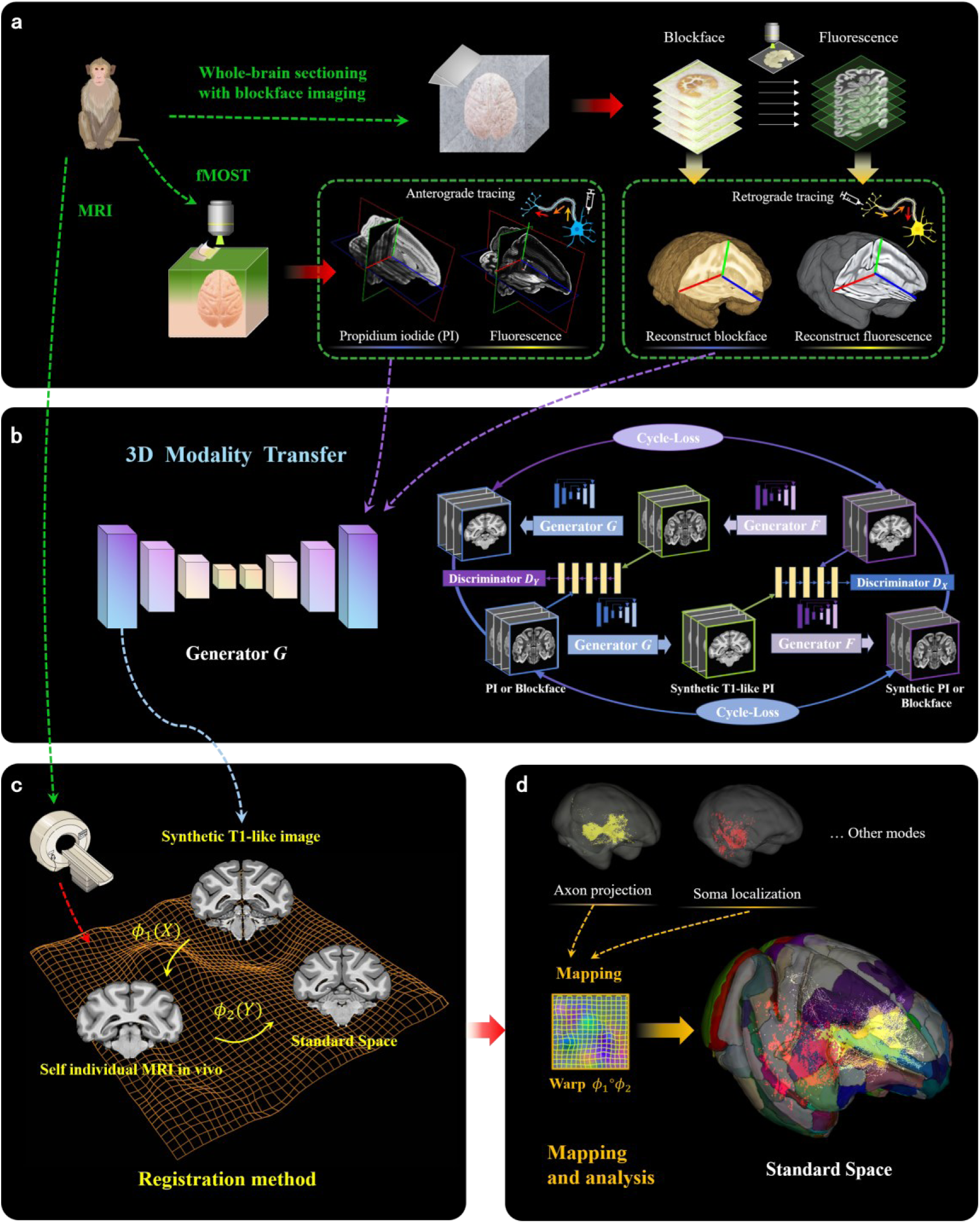
Overview of Macaca-star. **a**, Acquisition methods and 3D reconstruction of fMOST, Fluorescently Labeled Sections, and blockface Images. The fMOST separately acquired the PI channel and the GFP channel. Fluorescently labeled sections were used to obtain the stained-specific channel. **b, Whole-brain 3D modality transfer.** The 3D CycleGAN was utilized to transform fMOST PI and blockface images into the synthetic MRI T1-weighted modality. **c, In vivo self-individual MRI guided Whole-brain 3D registration.** The synthesized T1-like image, obtained through modality transfer from fMOST PI or blockface, was registered separately with the self-individual MRI and the standard template (NMT). **d, Mapping and analysis.** The data related to axon projections, soma spatial locations, or other modalities is mapping into the NMT standard space, and statistical analyses are performed using the atlas on the standard template.

The core objective of the preprocessing pipeline is to generate high-quality, smooth 3D optical images. For fMOST PI images, the primary preprocessing step involves the removal of stripe artifacts (Extended Data Fig. 1). In the case of block-face images, the main processing steps include brain segmentation and inter-slice alignment (Extended Data Fig. 2). Following preprocessing, we employ a 3D style transfer network to transform the smoothed PI or block-face images into MRI-modality-style images, thereby reducing modality-specific feature discrepancies relative to the NMT standard space images. To account for postmortem tissue distortions and inter-individual variability, we incorporate self-derived MRI as an intermediate registration bridge, ultimately generating a deformation field that maps 3D optical images to the MRI standard space (Fig. 1c). Notably, for 2D fluorescent slice images, we first perform 3D whole-brain registration of the block-face image to the standard space, followed by 2D registration between each fluorescent slice image and its corresponding block-face image (Extended Data Fig. 3). To address the substantial non-uniform deformations in 2D fluorescent slices and the modality differences between fluorescent and block-face images, we implement an integrated approach combining 2D style transfer learning with the Syn algorithm (Extended Data Fig. 3). Finally, we apply the deformation fields estimated from fMOST PI images or block face images to the fMOST GFP channel or thick fluorescent slice images of the same macaque brain, enabling precise mapping of axonal projections and neuronal soma distributions into the standard space (Fig. 1d).

### 3D Brain Modality Style Transfer

One of the key factors to improve the registration accuracy between cross-modal images is reducing the feature differences between the images. Although preprocessing provides us with smooth 3D optical images, there are still significant differences in brightness and texture between optical images and the MRI standard template. To achieve more robust and accurate registration of optical images to the standard space (e.g., the NMT standard space), we use an unsupervised 3D CycleGAN style transfer model^21^ (network structure shown in Extended Data Fig. 4), which converts optical slice images into a modality style closer to MRI. We chose to use a 3D model rather than a 2D model, which allows us to better leverage the 3D structural information of the images and ensure the consistency of brain tissue structure after transformation. We trained 3D style transfer models for both fMOST PI and Block face images respectively.

To validate the accuracy of the model, we performed a correlation analysis between the preprocessed images, style-transferred images, and self-individual MRI images with the NMT MRI template images. Whether fMOST PI images or Block face images, after style transfer network learning, the generated MRI-like images significantly improve in similarity to the NMT standard space images, closely resembling the self-individual MRI level (Fig. 2). For fMOST PI images, before and after style transfer network processing, the Pearson correlation coefficient with the NMT template significantly increased from -0.33 to 0.85, while the coefficient of determination (R-squared, R²) improved from 0.11 to 0.72 (Fig. 2a). For Block face images, the Pearson correlation coefficient increased significantly from 0.73 to 0.90, and the R² also improved from 0.53 to 0.81 (Fig. 2b).

**Fig. 2.**
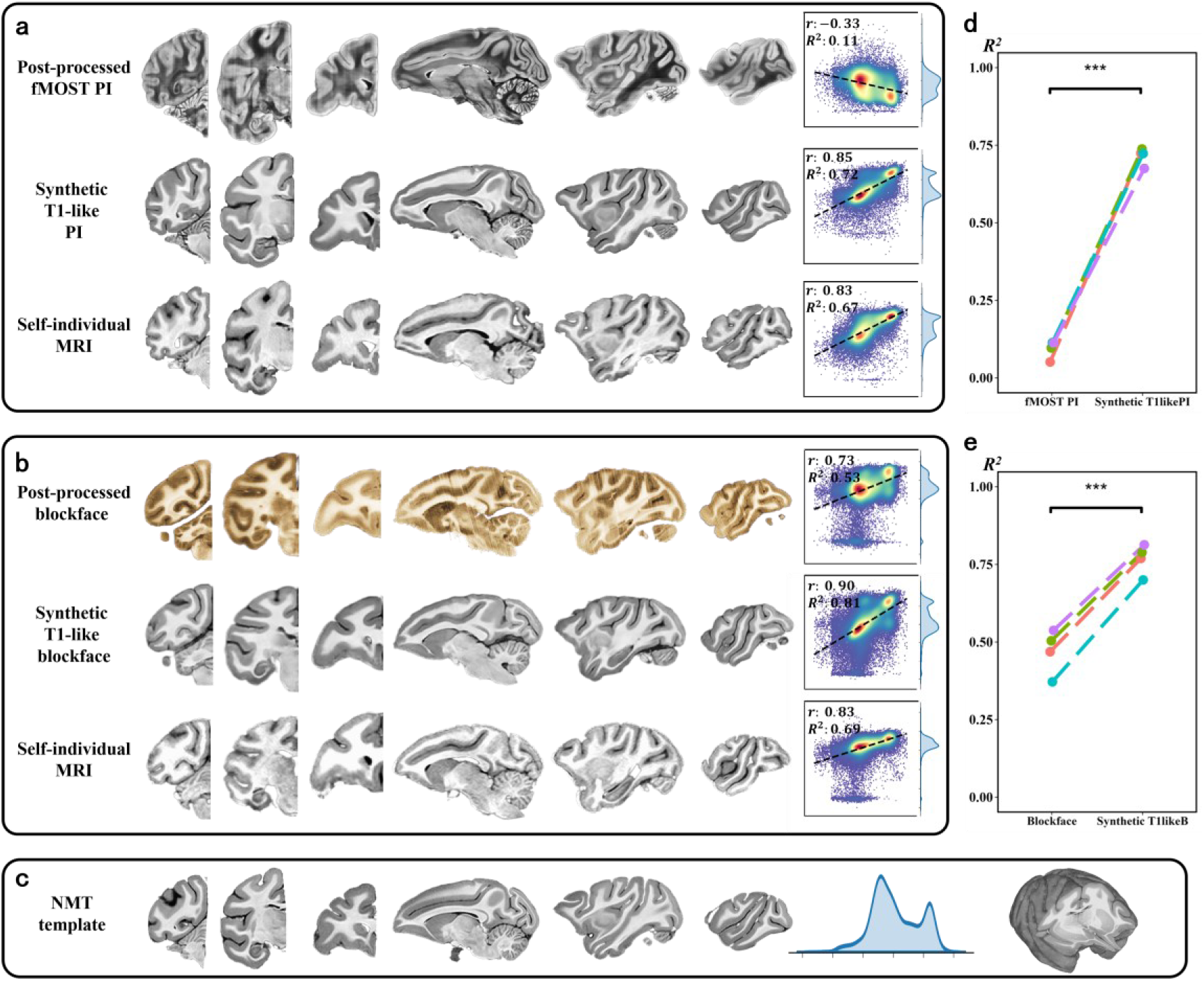
The similarity between different modalities and the NMT. **a**, The first, second, and third rows show slices in the coronal and horizontal planes for the post-processed fMOST PI images, the synthesized T1-like PI images following modality transfer, and the self-individual MRI images, respectively. On the right side of each row, the Pearson correlation coefficient (r) and the coefficient of determination (R²) between each image and the NMT standard template are displayed. **b,** The slices of the processed blockface images, their modality-transferred synthetic T1-like images, and self-individual MRI images are displayed. **c,** Visualization of different slices from the standard NMT template, along with their corresponding intensity distribution histogram. **d,** R ² were calculated for multiple sets of fMOST PI and their corresponding Synthetic T1-like PI images following modality transfer, comparing them with the NMT template. Paired t-tests were performed, revealing highly significant differences with p-values less than 0.001. **e,** Similarly, R² values were calculated for blockface images and their corresponding Synthetic T1-like Blockface images, with a p-value less than 0.001.

Furthermore, to assess the generalizability of the model, we applied the pre-trained style transfer models for PI and Block face images to other macaque brain samples. Both PI and block-face images showed significantly improved similarity to the NMI MRI template after processing through the trained 3D cycle-GAN model (Fig. 2d and 2e). This indicates that our trained model can be directly used for other ex vivo slice data collected using similar methods.

### fMOST-PI Registration accuracy

The most important aspect of cross-modal image registration is the integration of spatial location information. To better visualize and quantitatively evaluate the accuracy of the registration results, we display them in the individual space. We mapped the D99 cortical parcellation and the SARM subcortical nucleus parcellation, as defined in the NMT standard space, onto the 50×50×50 μm^3^ resolution fMOST PI individual space images for visualization and evaluation (Fig. 3a and video). The results of PI image registration to the NMT standard space are presented in the supplementary materials (Extended Data Fig. 5a). Both the D99 and SARM atlases were successfully mapped to the individual space (Fig. 3a). We chose five slices at the cortical and subcortical nuclear for display (Fig. 3a).

**Fig. 3.**
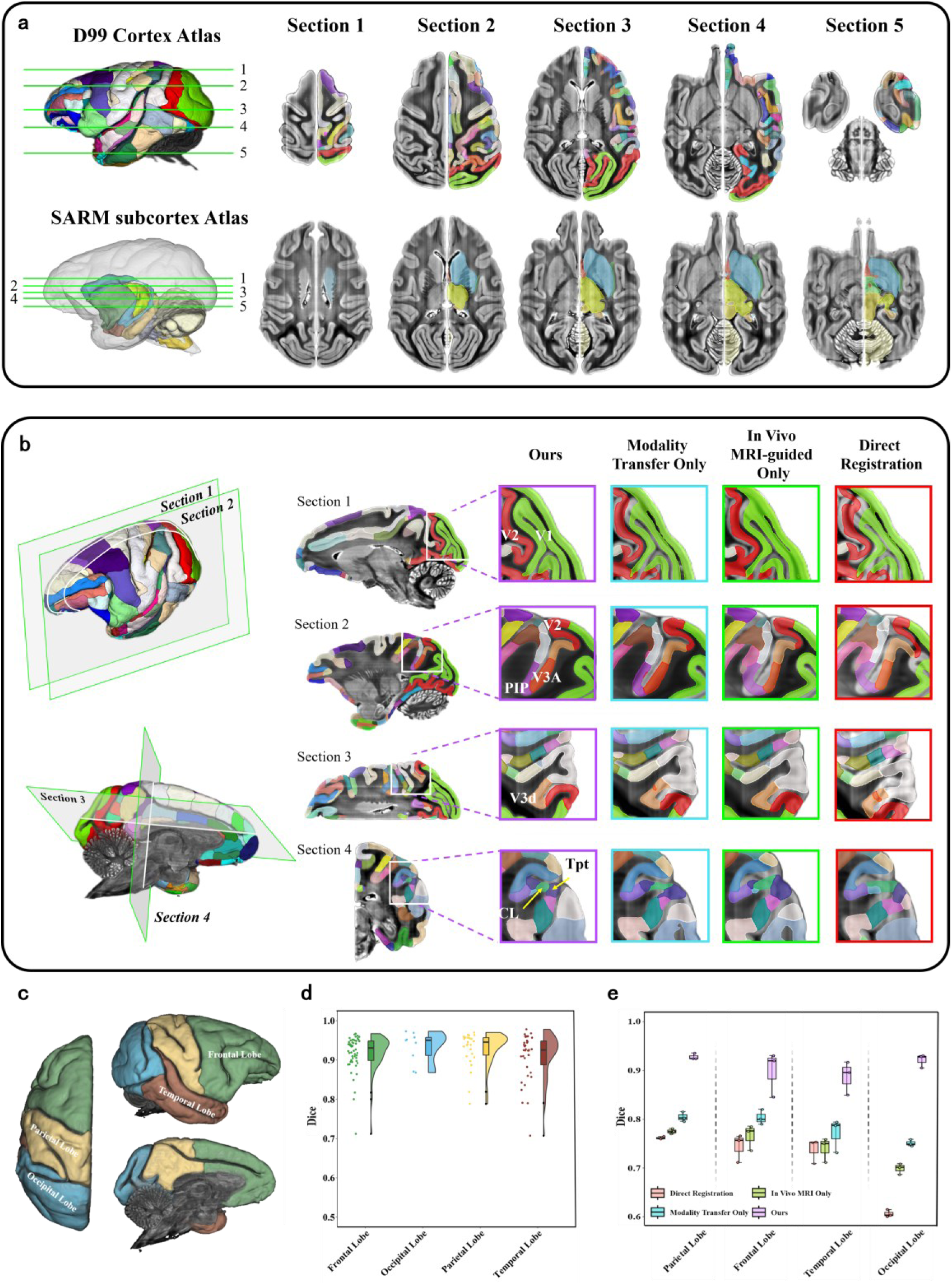
Evaluation of the 3D registration accuracy for fMOST PI image. **a,** The first and second rows respectively show the 3D registration results of the D99 atlas on the cortex and the SARM atlas on the nuclei, both registered to the fMOST PI, along with their corresponding slice registrations. **b,** Comparison of whole-brain registration methods for the fMOST PI. This figure compares the results of other three approaches: modality transfer only, in vivo self-individual MRI-guided registration only, and direct registration of the fMOST PI with the NMT template. The left side shows the spatial location of the displayed slices in the individual 3D space, while the right side presents the registration results for each method on the slices. **c,** The four major brain regions, frontal, occipital, parietal, and temporal lobes, registered to the fMOST PI are used to quantitatively assess registration accuracy. **d,** Quantitative evaluation of the whole-brain registration accuracy was performed by calculating Dice score for D99 brain regions within the frontal, occipital, parietal, and temporal lobes on a fMOST PI brain. **e,** The registration accuracy of different methods was quantitatively assessed by calculating the median Dice scores for subregions within the frontal, occipital, parietal, and temporal lobes, using three manually annotated D99 atlas on a macaque brain. Box plot: center line, median; box limits, upper and lower quartiles; whiskers, 1.5× interquartile range; points, outliers.

We compared results obtained by different registration strategies: 1) only style-transferred PI images without self-individual in vivo MRI; 2) only self-individual in vivo MRI without style-transferred PI; and 3) performing direct affine+Syn registration. The results show that only the method combining style transfer with MRI-guided registration achieves a good alignment of the PI images with the standard space. Omitting either condition results in significant misalignment for multiple subregion locations (Fig. 3b).

Finally, we invited three neuroanatomy experts to independently annotate the individual space PI images using the D99 atlas as a reference, which served as the gold standard for quantitatively evaluating the accuracy of the registration results. Based on this, we used the Dice coefficient to evaluate the consistency between the D99 atlas mapped from standard space to individual space at 50-micron resolution and the expert anatomical annotations. To facilitate better comparison of results, we consolidated multiple small subregions in the D99 atlas, ultimately reducing the number of subregions from 197 to 184, with specific consolidation details provided in the supplementary materials (Extended Data Table1). The consolidated whole-brain subregions demonstrated high registration accuracy, with median Dice coefficients of 0.922 [0.889, 0.941] (median [interquartile range]). Regional analysis revealed consistent performance across major lobar divisions (Fig. 3c): frontal lobe (0.919 [0.888, 0.936]), occipital lobe (0.934 [0.920, 0.941]), parietal lobe (0.925 [0.906, 0.946]), and temporal lobe (0.908 [0.863, 0.939]) (Fig. 3d). Comparative analysis showed that our method achieved median Dice scores approaching 0.9 across all major lobar regions, representing a 15-54% improvement in average Dice scores over alternative registration strategies (Fig. 3e).

### Fluorescence slices Registration accuracy

Traditional 2D histological slice images often exhibit significant irregular distortions, while block-face imaging maintains the brain’s structural morphology before tissue cutting. Therefore, for block-face images, following the same registration pipeline used for PI data, we successfully achieved accurate registration between block-face images and the MRI standard space. Similarly, we mapped both the D99 cortical parcellation and SARM subcortical parcellation onto individual block-face space images for visualization (Fig. 4a). To further demonstrate the robustness of our method, registration results from two other macaques are presented in the supplementary materials (Extended Data Fig. 6). Excitingly, the results demonstrate that our method can effectively register data with larger deformations (Extended Data Fig. 6b). Utilizing block-face registered images, we accomplished the alignment of significantly distorted 2D section images with the MRI standard space. The mapping of D99 cortical parcellation onto 2D fluorescent slice images is shown in Fig. 4b.

**Fig. 4.**
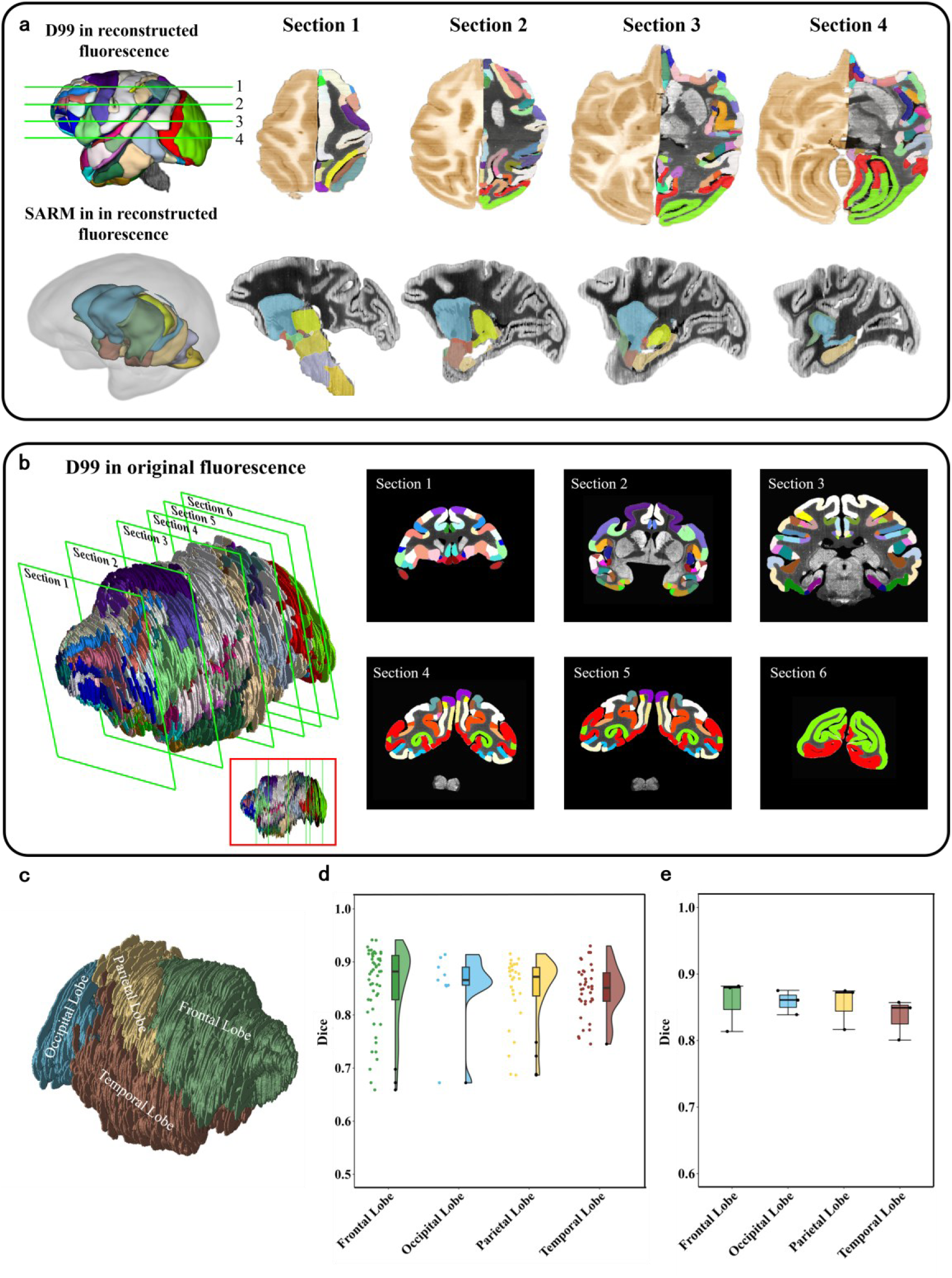
The registration accuracy of fluorescently labeled sections. **a,** The first row presents the registration results of the D99 atlas on the cortex. On the left is the registration of D99 in the 3D-reconstructed blockface space, while on the right are the registration results on sections at different levels. For each section, the left side displays the 3D-reconstructed blockface image, and the right side shows the 3D-reconstructed fluorescent image. The second row shows the registration results of the 3D-reconstructed fluorescent image using the SARM atlas for the nuclei. **b,** To satisfy the requirements for in situ analysis and quantitative evaluation of registration accuracy, we mapped the D99 atlas onto the original fluorescent slices. On the left, the result of D99 mapping onto the stacked original fluorescent slices is shown, while the right side presents the corresponding section display results. **c,** The four brain lobes are registered to the original fluorescent slices to assess registration accuracy. **d,** Quantitative evaluation of the whole-brain registration accuracy was performed by calculating Dice score for D99 brain regions within the frontal, occipital, parietal, and temporal lobes on fluorescently labeled sections. **e,** Quantitative evaluation of the overall registration accuracy for the four lobes, assessed by the median Dice scores of subregions within these lobes across three macaque brains. Box plot: center line, median; box limits, upper and lower quartiles; whiskers, 1.5× interquartile range; points, outliers.

Similarly, we invited three anatomical experts to independently annotate the raw fluorescent slice data from three macaque brains using the D99 atlas as reference, serving as the gold standard to evaluate the accuracy of our registration. The parcellation results of all subregions in one macaque brain, divided according to four major lobes (Fig. 4c), are shown in Fig. 4d, while the results for the other two brains are provided in the supplementary materials. Quantitative analysis revealed consistent registration accuracy across all brains, with median Dice coefficients (interquartile ranges) for major lobar regions as follows: frontal lobe (0.880 [0.826, 0.913]), occipital lobe (0.875 [0.860, 0.885]), parietal lobe (0.874 [0.854, 0.890]), and temporal lobe (0.857 [0.839, 0.883]) (Fig 4.d). Across all three specimens, median Dice scores remained stable: frontal lobe (0.880), occipital lobe (0.861), parietal lobe (0.872), and temporal lobe (0.849) (Fig. 4e).

### Macaque brain connectivity profile

We applied Macaca-Star to integrate and analyze anterograde and retrograde tracing connectivity results from different imaging techniques in standard space. Among these, fMOST technology, with its remarkable advantages in microscopic imaging at single-cell resolution, has become a crucial imaging modality for acquiring complete projection pathways of individual neurons. The fMOST data acquisition system supports multi-channel synchronous acquisition, enabling simultaneous collection of PI images reflecting architectural information and GFP channel data characterizing neuronal projection features. By registering fMOST PI data to the NMT standard space, we obtained the deformation field from individual space to standard space, thereby achieving the reconstruction and mapping of neuronal projection profile acquired from the GFP channel into standard space. We demonstrate the whole-brain projection patterns resulting from AAV virus injection in the substantia nigra; however, due to viral leakage into other brain regions during injection in the samples used, this study provides only qualitative demonstrations of projection data in original individual space and their mapping results in standard space (Fig. 5a). After data processing and registration processes, we achieved both quantitative analysis and visualization of axonal projections in standard space, along with reverse mapping of standard space atlases (e.g., D99) to individual space, facilitating accurate spatial localization and quantitative assessment of cortical projection patterns.

**Fig. 5.**
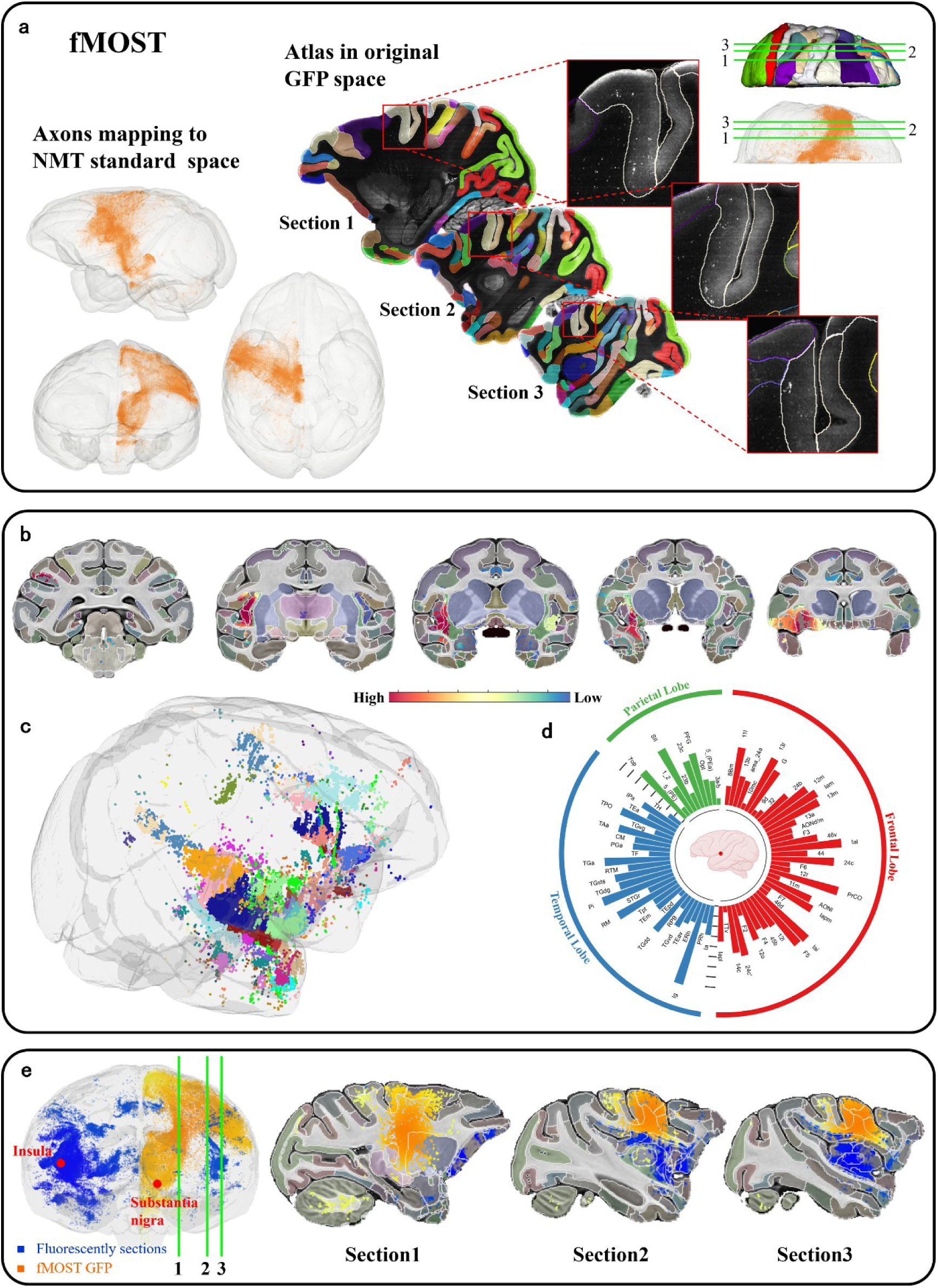
Whole-brain axonal projections and soma distribution architecture, and regional analysis in standard space. **a,** The axonal projections can be presented and analyzed in two ways. On the left, the axons are mapped to standard space using the Macaca-star registration method, enabling standardized presentation and analysis. On the right, the D99 atlas is mapped to the individual fMOST space for in situ analysis, preserving the specificity within the individual anatomical structure. (The injection site is the substantia nigra) **b,** The soma locations reconstructed from fluorescently labeled sections of a macaque brain are mapped to the NMT standard template through the Macaca-star pipeline. The first rows display the distribution of soma on the coronal planes of the NMT template. **c,** The soma locations are mapped to standard space, with soma distributions in different brain regions color-coded according to the D99 atlas. **d,** Additionally, soma distribution statistics were performed based on the hierarchical regions of the D99 atlas (The injection site is the insula). **e,** The soma distribution derived from retrograde-labeled fluorescent sections and the neuronal projections traced by anterograde fMOST GFP were mapped into a common standard space. The injection sites were located in the Insula and Substantia Nigra, respectively. The right panel displays sagittal slices, with corresponding locations indicated on the left image.

Following registration to the NMT template, the distribution of retrogradely labeled neurons from ID subregion in insula cortex was visualized (Fig. 5b). Labeled neurons were predominantly localized to the temporal lobe, frontal lobe (including prefrontal, frontal, and cingulate regions), parietal lobe, and several subcortical areas (e.g., amygdala and thalamus) (Fig. 5b). Spatial analysis revealed distinct regional specificity: within the temporal lobe, labeled neurons were concentrated in Ig, Pi, and RM, regions proximal to the injection site. In the frontal lobe, labeling was primarily observed in ventral (areas 11l, 12m, 13l, 13m, lam, lai, PrCO, F5) and medial (areas 24c, 24c’) regions. Similarly, in the parietal lobe, labeled neurons were predominantly distributed in ventral (SII, 7op, PFG) and medial (23c) areas. These findings align with previous individual space analyses, further validating the accuracy of our registration approach. To further demonstrate the robustness of our method, we also registered fluorescence slice data from two additional channels of the same macaque into the standard space (Extended Data Fig. 7). Finally, the results demonstrate that our method enables the integration of both anterograde and retrograde tracing results obtained from different samples and modalities into standard space (Fig. 5e).

## Discussion

We have established a fully automated pipeline that integrates deep learning with in vivo self-individual MRI-guided registration to register macaque brain optical slices to the NMT MRI standard space. This registration pipeline enables the mapping and integration of Cytoarchitecture, projection profile, soma location, and other data from various individual spaces into the NMT standard space. There have been studies on mice that achieved 2D slice registration, without guidance from in-vivo individual image, to the standard space using only image style transfer^12^, while on marmosets that performed MRI-guided registration alone without image style transfer^17^. However, our results indicate that either style transfer or MRI-guided registration themself cannot effectively achieve precise registration of macaque 2D optical slices to the MRI standard space (Fig 3 and Extended Data Fig. 6). This could be due to the greater ex vivo deformation of macaque optical slices, along with larger individual differences among macaques.

Macaca-Star can be easily applied to different samples from various data sources. We successfully tested the integration of anterograde tracing data from fMOST (Fig 5.a) and retrograde tracing data from optical slices into the standard space (Fig 5.b). Furthermore, the trained 3D style transfer model exhibits strong generalizability (Fig 2), allowing direct application to new fMOST PI data or block-face data without requiring additional training. Additionally, one of the advantages of our method is its ability not only can individual optical images be registered to standard space, but standard space-defined subregion parcellations can also be precisely mapped back to individual space. This feature makes it particularly suitable for applications requiring preservation of in situ spatial information, such as single-neuron reconstruction from fMOST data^23^ or spatial transcriptomics^9^ analyses.

Although our method requires multiple types of data, these are routinely collected in many studies. For example, in vivo MRI acquisition has been widely used for spatial localization of viral injections and staining labels in large animals^23–25^. The acquisition of Block face images is also commonly used to reconstruct atlases of various species^6, 26, 27^. MOST facilitates the acquisition of information from diverse species and multiple staining methods^23, 28–31^. Therefore, our workflow could be expanded in the future to register additional optical modalities, such as Spatial Transcriptomics^9^, from various species, including the marmoset^32^ and human^6^, to the standard space.

Macaca-Star is primarily focused on the registration of fine cortical subregions. For some fine subregions of subcortical nuclei^33, 34^, their boundaries can be easily distinguished in optical slices. However, due to the resolution of the NMT standard space that it is constructed using MRI, it cannot distinguish the boundaries well, which makes it difficult to achieve precise subregion registration of subcortical nuclei in the stand space. In the future, the creation of macaque templates based on optical images, or higher-resolution atlases, will address the challenge of registering fine subregions of subcortical nuclei. Finally, Macaca-Star is an open-source, multi-module design that is scalable for preprocessing multiple modal slice data and integrating cross-modal and cross-individual information.

## Acknowledgments

This work was partially supported by the National Key R&D Program of China (Grant Nos. 2022YEF0203200 and 2022YFA1603604); the STI2030-Major Projects (2021ZD0200104); the National Natural Science Foundation of China (Grant Nos. 82260227, 61890950, and 62401185); Hainan University Research Start-up Fund (KYQD(ZR)20072 and KYQD(ZR)22074); the PhD Scientific Research and Innovation Foundation of The Education Department of Hainan Province Joint Project of Sanya Yazhou Bay Science and Technology City (HSPHDSRF-2024-08-010).

## Methods

### Animal care and use

All experiments related to use of macaque from the fMOST group, followed procedures approved by the Institutional Animal Ethics Committee of Huazhong University of Science and Technology (approval No. S166). All the experiments carried out in IoN were approved by the Ethics Committee of the Institute of Neuroscience, Chinese Academy of Science (ION-2019009 and CEBSIT-2020022).

### Data preparation

The NMT v2 standard template, D99 atlas, and SARM atlas were obtained from AFNI (https://afni.nimh.nih.gov/pub/dist/doc/htmldoc/nonhuman/macaque_tempatl/template_n mtv2.html). The NMT template represents a high-resolution in vivo MRI macaque brain standard with an isotropic voxel size of 250 μm^35^. This comprehensive template provides both precise tissue segmentation (gray matter, white matter, and CSF) and an integrated 3D digital D99 atlas derived from 2D histological sections. The original D99 atlas contains 197 subregions^34, 36^. To facilitate better quantitative comparison of registration results, we merged multiple very small subregions in the D99 atlas (The details showed in the Extended Data Table1). The finally D99 atlas used for our analysis contains 184 subregions. Additionally, the SARM^37^ subcortical atlas, which was constructed from high-resolution ex vivo structural imaging data and has been registered to the NMT standard template. The SARM atlas features multiple spatial hierarchical scales; for the current study, we utilized the second hierarchical level for all analyses.

fMOST dataset includes brain imaging data from four macaques, encompassing two main imaging channels: the PI channel and GFP channel. Just one macaque underwent MRI scanning to acquire T1-weighted T1w images (TE = 3.6 ms, TR = 10.56ms, FA = 10 degree, number of average = 3), with an isotropic resolution of 0.5mm. For GFP imaging, we labeled the macaque brains using AAV anterograde tracing technology. A detailed description of the sample preparation and imaging methods has been provided in previous studies^38^.

In this study, retrograde tracing experiments were performed in four adult cynomolgus monkeys (Macaca fascicularis). Before the injection experiment, each underwent MRI scanning to acquire T1-weighted T1w images (TE = 2.8 ms, TR = 2200 ms, FA = 9 degree, number of average = 4), with an isotropic resolution of 0.5mm. A detailed description of the sample preparation and imaging methods has been provided in previous studies^39^.

### fMOST preprocessing

#### Removing striping artifacts

To robustly and quickly remove artifacts at a resolution of 50×50×50μm³, we employed a method based on fitting the center of the focal plane (Extended Data Fig. 1a). The specific steps are as follows: (1) Downsample the original resolution to 50*50*50μm³ and reconstruct the 2D slices into 3D. (2) Project the entire reconstructed 3D image onto the slice plane and average the projection to obtain the average projection image. (3) Select the direction perpendicular to the stripe artifact, then re-project and average the projection along this direction to obtain the intensity curve C(x) along the edge. (4) Calculate the local maxima of the one-dimensional signal by setting the minimum peak height, width, and the minimum distance between adjacent peaks, identifying each vertex of the curve. (5) Fit the obtained vertices using a Bessel function to derive the fitted curve C_b_(x). (6) Divide the obtained curve C_b_(x) by the the intensity curve C(x), thus obtaining the correction function ratio(x) = C_b_(x)⁄C(x). (7) Use the correction function to restore the image. Repeat the above steps for the artifact on the other side of the image.

#### Nonuniform intensity correction

We first performed denoising on the 50×50×50μm³ PI images using a Gaussian filter with a σ value of 1. Then, we mapped the images to a reference space with a resolution of 250×250×250μm³ using affine transformation, where the reference was the MRI images of the individual macaque brain. To correct for intensity non-uniformity, we applied N4BiasFieldCorrection^40^ from ANTs, which helps reduce the impact of light source intensity variations on the image. Afterward, we enhanced the corrected images using CLAHE, significantly improving the local contrast of the images.

### Preprocessing for 2D fluorescent sections

#### Segmentation of the Consecutive blockface images

Before performing tissue section imaging, we first acquired the corresponding blockface images for each slice, which preserved the morphological structure prior to tissue slicing. However, blockface images contain significant background information. To handle this background, we used the SAM2 (Segment Anything 2)^41^ model for segmentation of the Consecutive blockface images (Extended Data Fig. 2a). SAM2 consists of five modules: image encoder, prompt encoder, mask decoder, memory attention, and memory bank. The memory attention is to condition the features of the current frame with those of previous frames, predictions, and any new prompts. The memory bank is responsible for maintaining predictions of neighboring frames, including past target predictions and stored prompts. Since the blockface images we captured are consecutive and the adjacent slices maintain similar sizes and structures, we perform segmentation predictions by treating the consecutive images as frames in a continuous video.

We initialize SAM2 based on a pre-trained model and fine-tune it accordingly. Given that our data consists of consecutive static images, we designed a training strategy that involves randomly sampling the blockface dataset from macaque subjects. For each subject’s dataset, an alternating training strategy is used to randomly sample both images and videos for training. Additionally, we ensure that the set of images sampled from the static images for video training is continuous. The slices are also organized in descending order based on their size. The specific steps are as follows: First, we performed automatic segmentation without prompts on all blockface images. Then, we inspected the segmentation quality, and for images with segmentation errors, we corrected them by adding prompts to guide the model (Extended Data Fig. 2b).

#### 3D blockface reconstruction

Due to platform or tissue movement during the acquisition of blockface images, and since the macaque brain was divided into anterior and posterior halves along the coronal plane, there is an offset within the blockface slices, which prevents direct 3D reconstruction (Extended Data Fig. 2e). To achieve 3D reconstruction of the blockface, we first calculate the mean squared error (MSE) between slices to identify the specific locations of the offsets (Extended Data Fig. 2d), which can be formulated as:

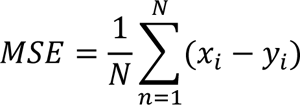

where *N* represents the number of voxels in the image, where *x* refers to the voxels on a slice, and *y* denotes the corresponding voxels on adjacent slices that are spatially aligned with *x*.

Once the offset locations are identified, we locate the center of the blockface slices and align the first offset slice with the adjacent slice. Then, the remaining offset slices are corrected by applying the same shift distance (Extended Data Fig. 2c). Due to the shift of the blockface slices relative to the camera, affine deformations occur. We perform an affine transformation on the first offset slice in relation to the adjacent slice. Similarly, the same affine matrix is applied to the remaining slices, and finally, 3D reconstruction is performed (Extended Data Fig. 2f).

#### 3D reconstruction of 2D fluorescent sections

We developed a 3D reconstruction method for fluorescence tissue sections derived from blockface imaging (Extended Data Fig. 3a). Initially, we employed a 2D CycleGAN model to transform the fluorescence images of 2D sections into a blockface modality, ensuring modality consistency. Subsequently, we performed affine registration of the synthetic blockface-like sections with their corresponding blockface images to align the overall tissue slices. Following this, we applied a watershed-based segmentation method to accurately segment the slices. Specifically, we segmented both the fluorescence and corresponding blockface slices, with a particular focus on regions exhibiting significant deformation, such as the separated left and right brain tissue and the temporal lobe. After segmentation, we aligned the tissue regions in the fluorescence image to their corresponding regions in the blockface image using the synthetic blockface-like sections, thereby restoring severely misaligned tissue structures in the fluorescence image to their correct anatomical positions (Extended Data Fig. 3b). Throughout these steps, we computed affine matrices and deformation fields, which were subsequently used to map the atlas to the individual’s original fluorescence slices and to map soma locations from the original slices into standard space.

#### 3D modality transfer of macaque brain images

In this study, we utilized a 3D cross-modal transfer network to convert cross-modal registration into intra-modal registration (Fig. 1c). The network employed is the Cycle-Consistent Adversarial Network (CycleGAN), which enables the conversion of fMOST PI or 3D-reconstructed blockface images into the MRI T1-w modality. The key difference from the original CycleGAN is that we employ a 3D approach instead of a 2D one. This allows us to fully leverage the three-dimensional structural information of the images, ensuring consistency in histological and anatomical structures. The 3D CycleGAN is a model with two directions: forward cycle and backward cycle. In the forward cycle, there are two mappings involved: from domain *X* to *Y*, *G*: *X* → *Y*, and then from the generated domain Y back to domain *X*, *F*: *Y* → *X*. The backward cycle is the reverse of this process. The main objective is to transform fMOST PI or 3D-reconstructed blockface into the

T1-weighted (T1w) modality of MRI, with particular focus on the forward process, especially the generator *G* (Extended Data Fig. 4b). The generator *G* is primarily composed of 9 ResNet blocks (Extended Data Fig. 4d). Additionally, the model includes two adversarial discriminators (Extended Data Fig. 4a), *D*_*X*_ and *D*_*Y*_. *D*_*Y*_ is primarily used to differentiate images from domain *X* and the synthetic images in domain *Y*, which correspond to real T1w images and synthetic T1-like fMOST PI or 3D-reconstructed blockface images. Meanwhile, *D*_*X*_ is responsible for distinguishing optical images. The discriminator *D* employs Binary Cross-Entropy (BCE) loss, which is formulated as follows:

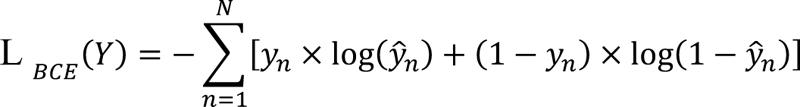

where *Ŷ* represents the model’s predictions, and y represents the ground-truth. The entire network’s loss consists of two components: adversarial losses and cycle consistency losses, which can be formulated as:

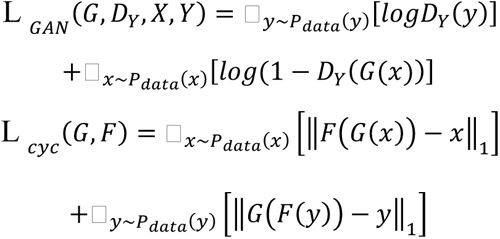

The full loss is:

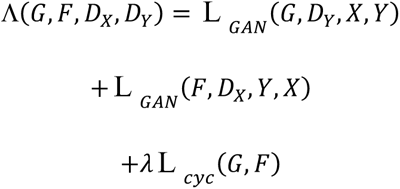

where λ controls the relative importance of the two losses.

We utilized two types of optical images: fMOST PI and 3D-reconstructed blockface. Therefore, we trained the model separately using each dataset with the NMT template. Initially, the model was trained with fMOST PI and the NMT template. To ensure spatial consistency, we aligned the fMOST PI and 3D-reconstructed blockface images to the NMT template, resulting in an image size of 256 × 312 × 200. During training, to accommodate memory constraints, we set the batch size to 1 and divided the images into 128 × 156 × 200 patches (Extended Data Fig. 4c) to fully leverage the 3D anatomical information. The learning rate was set to 0.0002, and the model was trained for a total of 250 epochs. Subsequently, we applied the same training strategy to blockface and NMT.

#### Preprocessing for self-individual MRI

We developed a preprocessing pipeline for macaque brain MRI images based on the HCP method for human brain MRI processing^42^ (Extended Data Fig. 8), in order to isolate non-brain tissues and extract brain tissue. The steps are as follows: (1) First, we corrected the orientation of the MRI images so that the individual’s MRI alignment matched the NMT standard template. (2) To minimize the interference of excess tissues during registration, we cropped the neck region. (3) We performed affine and nonlinear registration between the individual MRI and the standard template, obtaining the affine matrix and deformation field. (4) Using the inverse deformation field, we mapped the brain mask from the NMT standard space to the individual MRI space. (5) The brain tissue was then extracted using the mask. (6) We applied N4BiasFieldCorrection from ANTs to correct the bias field in the image. (7) A denoising method based on non-local means filtering was used to reduce noise in the low signal-to-noise ratio MRI images. (8) Finally, we further registered the synthesized image, now aligned to the MRI, to the NMT standard template. This process began with an affine transformation to globally align the synthesized image to the NMT standard template.

#### Synthetic T1-like images registration to NMT by self-MRI

We developed a registration method based on individual in vivo MRI images. First, using the skull-stripped MRI T1-w as a reference, we registered the T1-like PI or 3D-reconstructed blockface images synthesized with a 3D CycleGAN model. During the registration process, we first applied a global transformation to the synthetic images. By minimizing the global correlation (GC) metric, we performed an affine linear transformation to normalize the images to the same scale and resolution as the MRI. The GC metric can be formatted as:

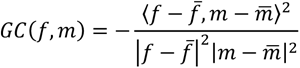

where *f* represents the vector of T1-w image pixel intensities, and m denotes the synthetic image. *f̄* and *m̄* are the mean values of *f* and *m*, respectively. The notation <,> represents the inner product. Subsequently, we employed the SyN method for nonlinear registration, using mutual information (MI) as the similarity criterion^43^, at a spatial size of 250 × 250 × 250μm³. This approach precisely registers the synthetic image to the individual MRI. The negative mutual information *S* between the T1w image, serving as the reference, and the transformed synthetic image is formulated as:

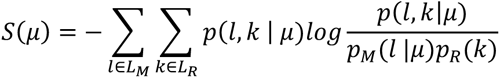

where μ is a vector of the transformation parameters. The joint probability distribution is denoted by *p*, while *p*_*M*_ and *p*_*R*_ represent the marginal probability distributions of the synthetic moving image and the T1w reference image, respectively.

Subsequently, we applied the same registration method and similarity criterion to align the T1-like PI or reconstructed blockface image, which had been registered to the T1w image, to the standard NMT template.

