## Supplemental files for "Fully Automated and Scalable Pipeline for Macaque Brain Registration"

**Supplementary Figures and table**

Supplementary Figures

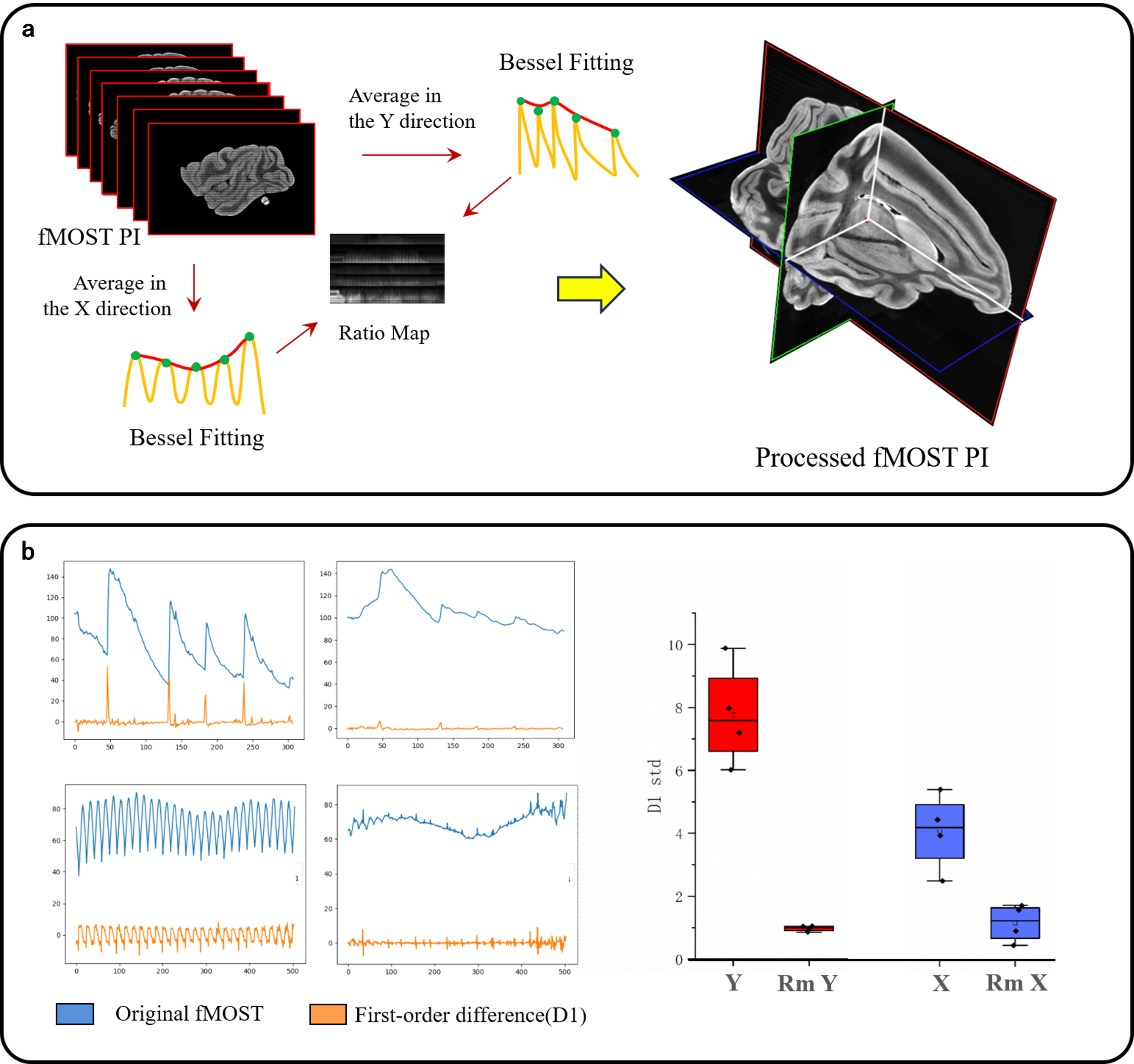

**Extended Data Fig. 1 | Removing striping artifacts. a,** Workflow for removing stripe artifacts from fMOST PI images. **b,** On the left side, the first and second rows show the intensity variations of the averaged posterior projections along the y and x axes for both the original and artifact-corrected fMOST PI images, followed by the calculation of their first-order differences. Additionally, the variance of the first-order difference along both directions was calculated using four fMOST PI datasets and displayed on the right side.

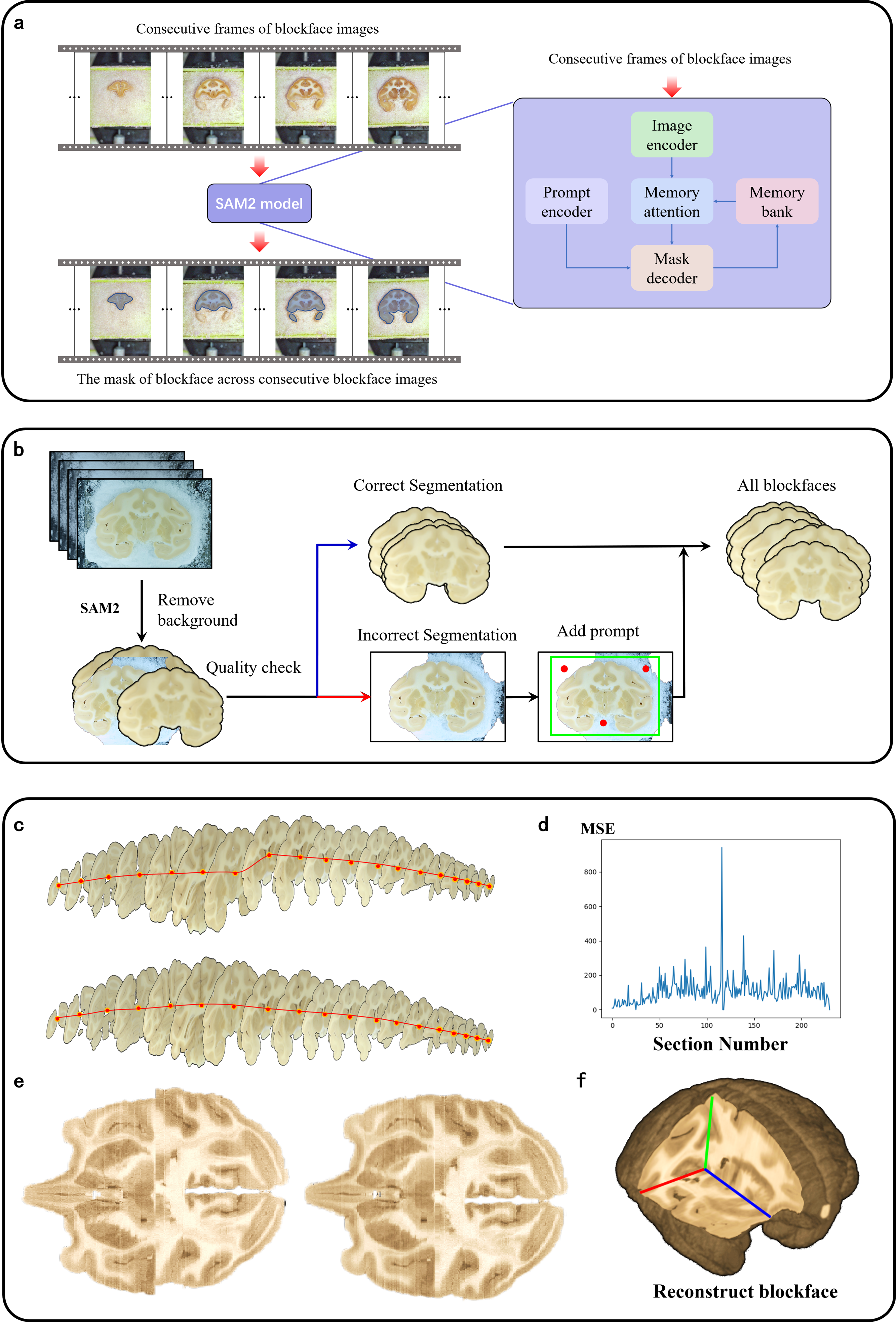

**Extended Data Fig. 2 | Brain extraction and 3D reconstruction from blockface images. a,** Brain extraction from blockface images and the architecture of the SAM2 model. **b,** Quality control of blockface image segmentation. **c,** Alignment of blockface centers. **d,** MSE between adjacent blockface image slices. **e,** Comparison of blockface images before and after 3D reconstruction. **f,** The image displays the final 3D reconstructed volume obtained from the original blockface slices.

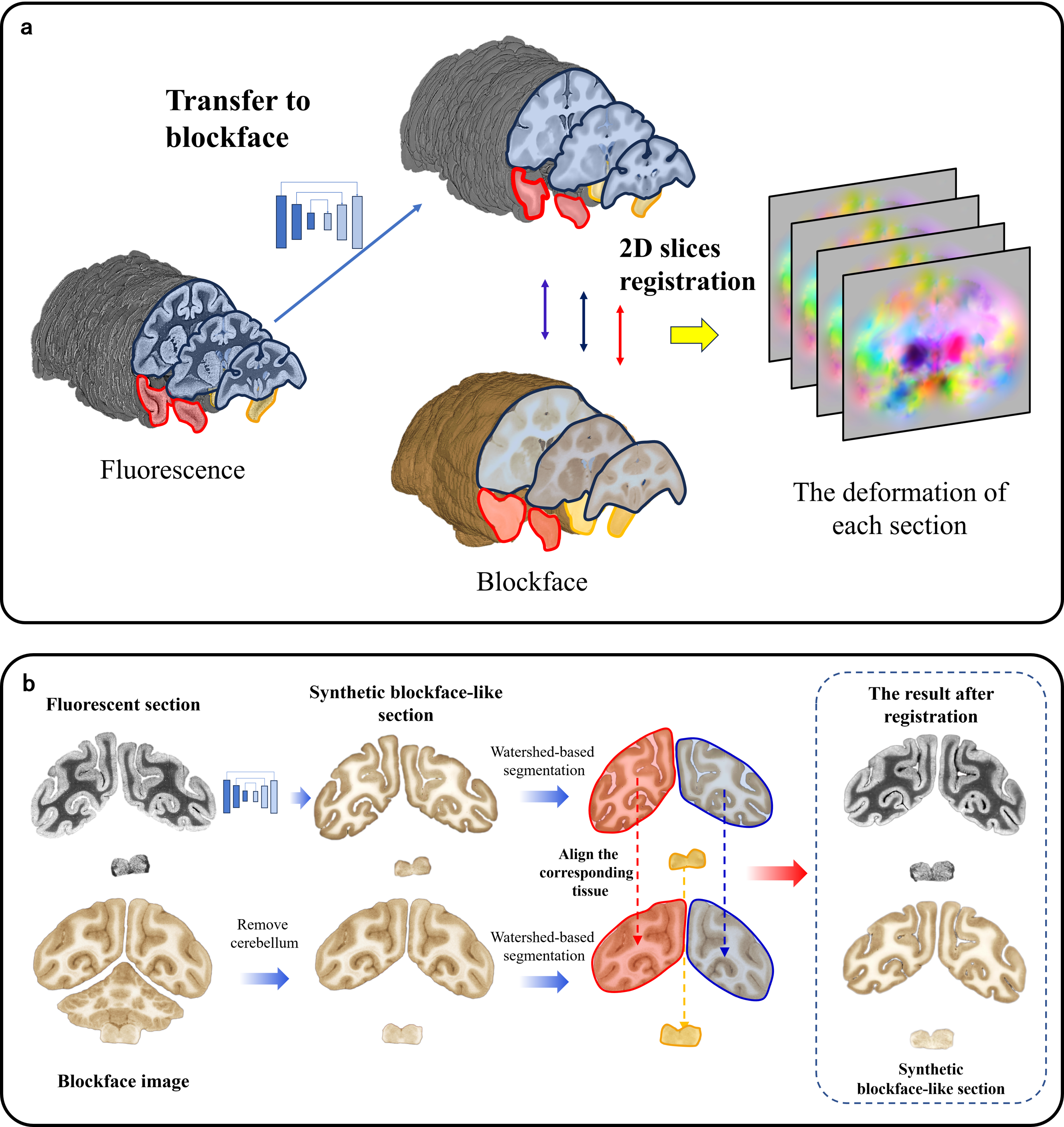

**Extended Data Fig. 3 | 3D reconstruction of 2D fluorescent sections. a,** Fluorescent sections aligned to the corresponding blockface images through the modality transfer model. **b,** After modality transfer of the fluorescent sections, tissue regions from different blocks of the fluorescent sections and their corresponding blockface images were segmented and further aligned to the same anatomical structures.

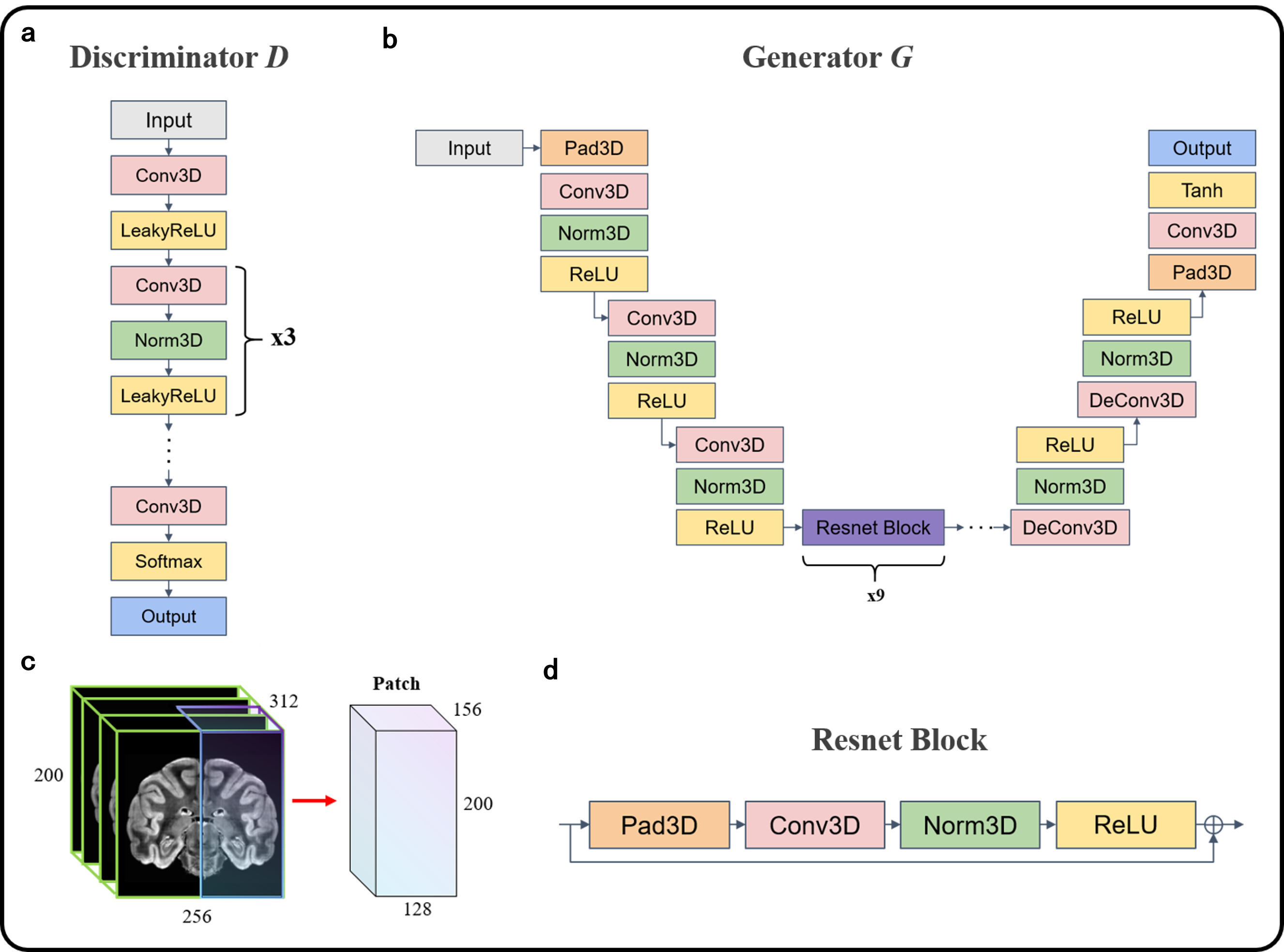

**Extended Data Fig. 4 | Architecture of the modality transfer deep learning network. a, b, d,** The specific architectures of the network's discriminator, generator, and ResNet block. **c,** The size of the input patch to the network.

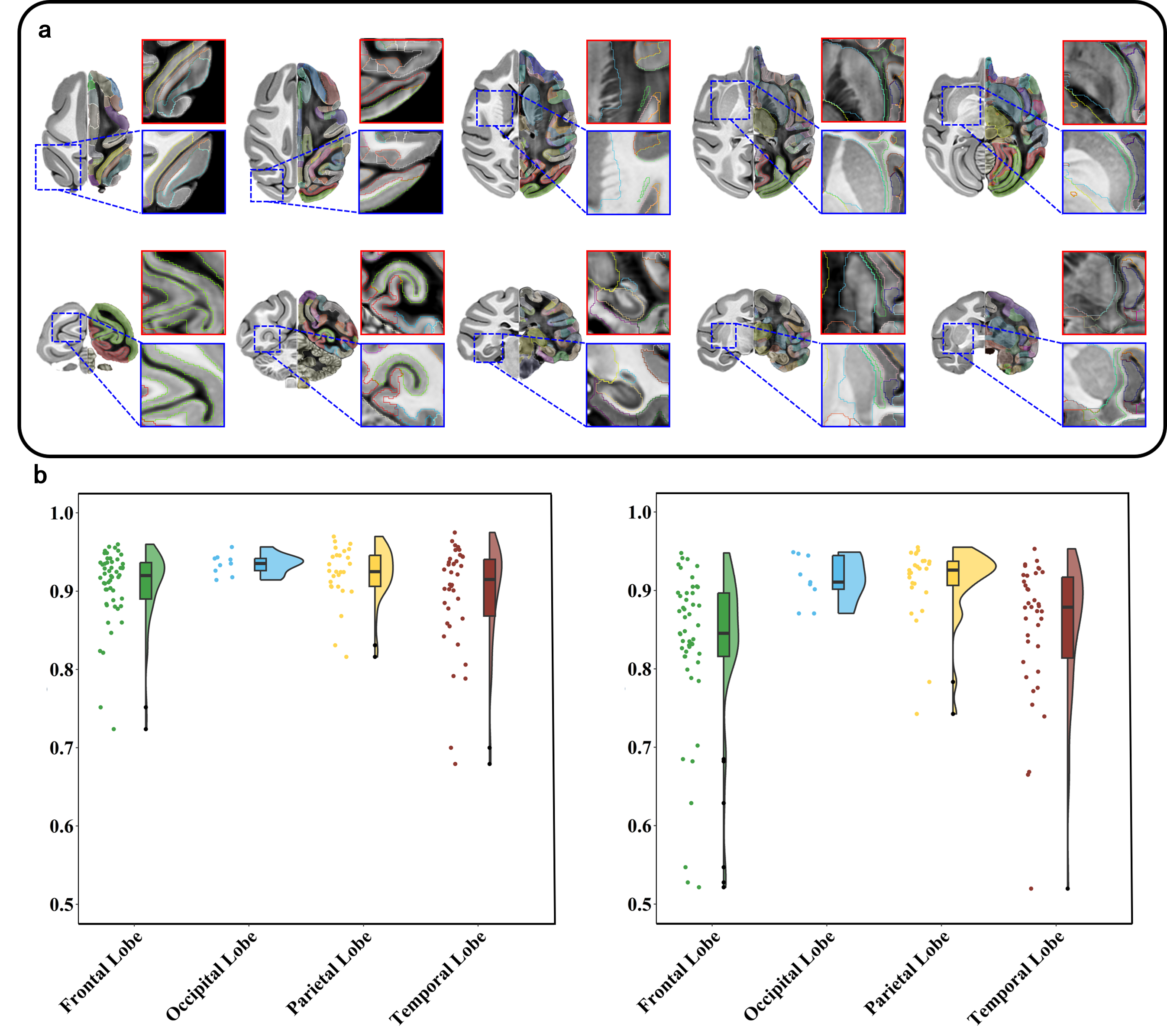

**Extended Data Fig. 5 | Qualitative and quantitative evaluation of the registration accuracy of the fMOST macaque brain image. a,** Results of the fMOST PI image registration to the NMT standard template. The left side of each brain shows the NMT standard template, while the right side displays the corresponding fMOST PI image at the same location. **b,** Distribution of Dice scores calculated from the D99 atlas annotations by two anatomical experts and the registration results across different brain lobes.

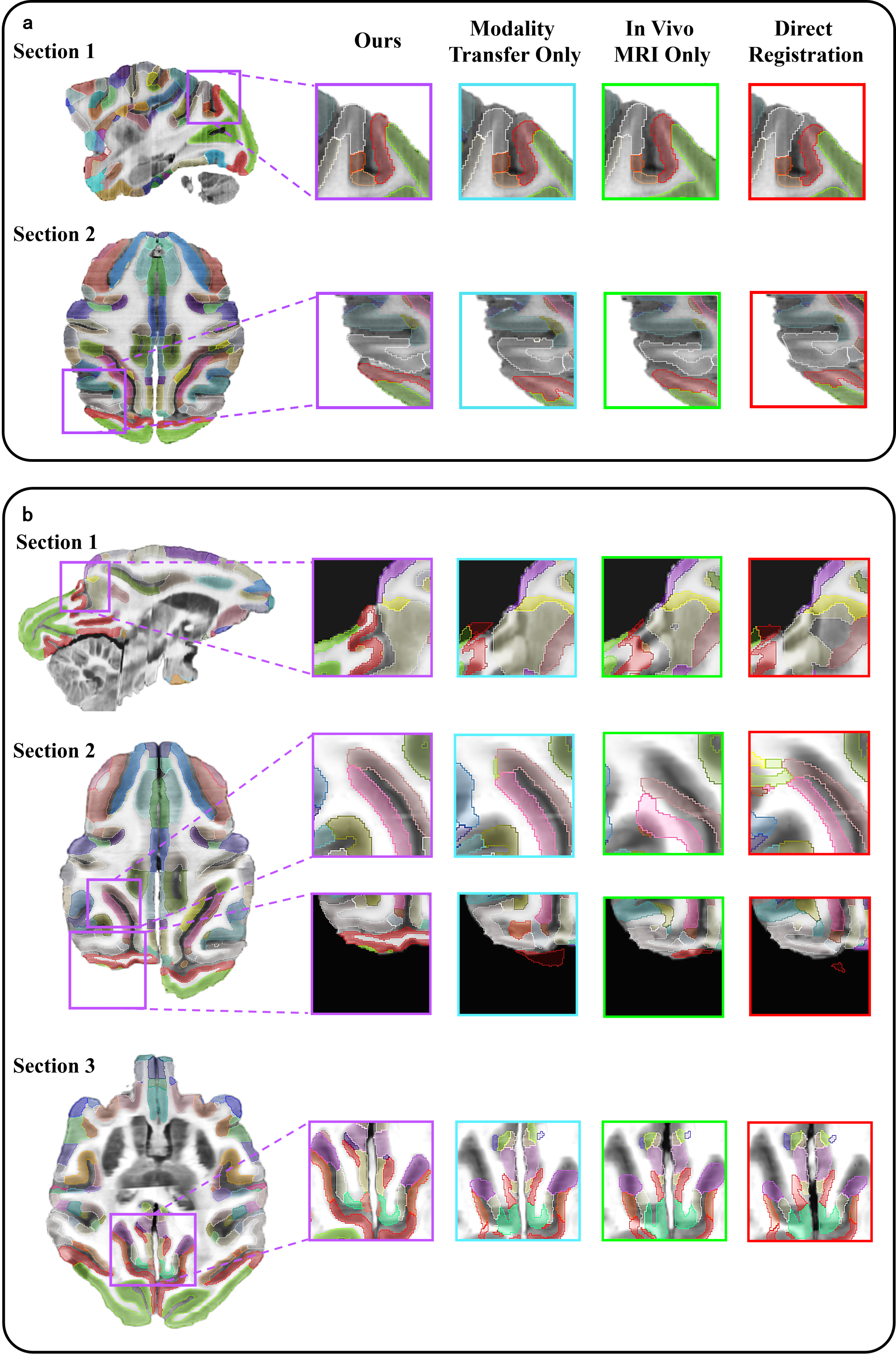

**Extended Data Fig. 6 | Comparison of registration results for 3D-reconstructed blockface brain images using different methods. a** and **b** show the results for two different macaque brains, with colored boxes representing different methods.

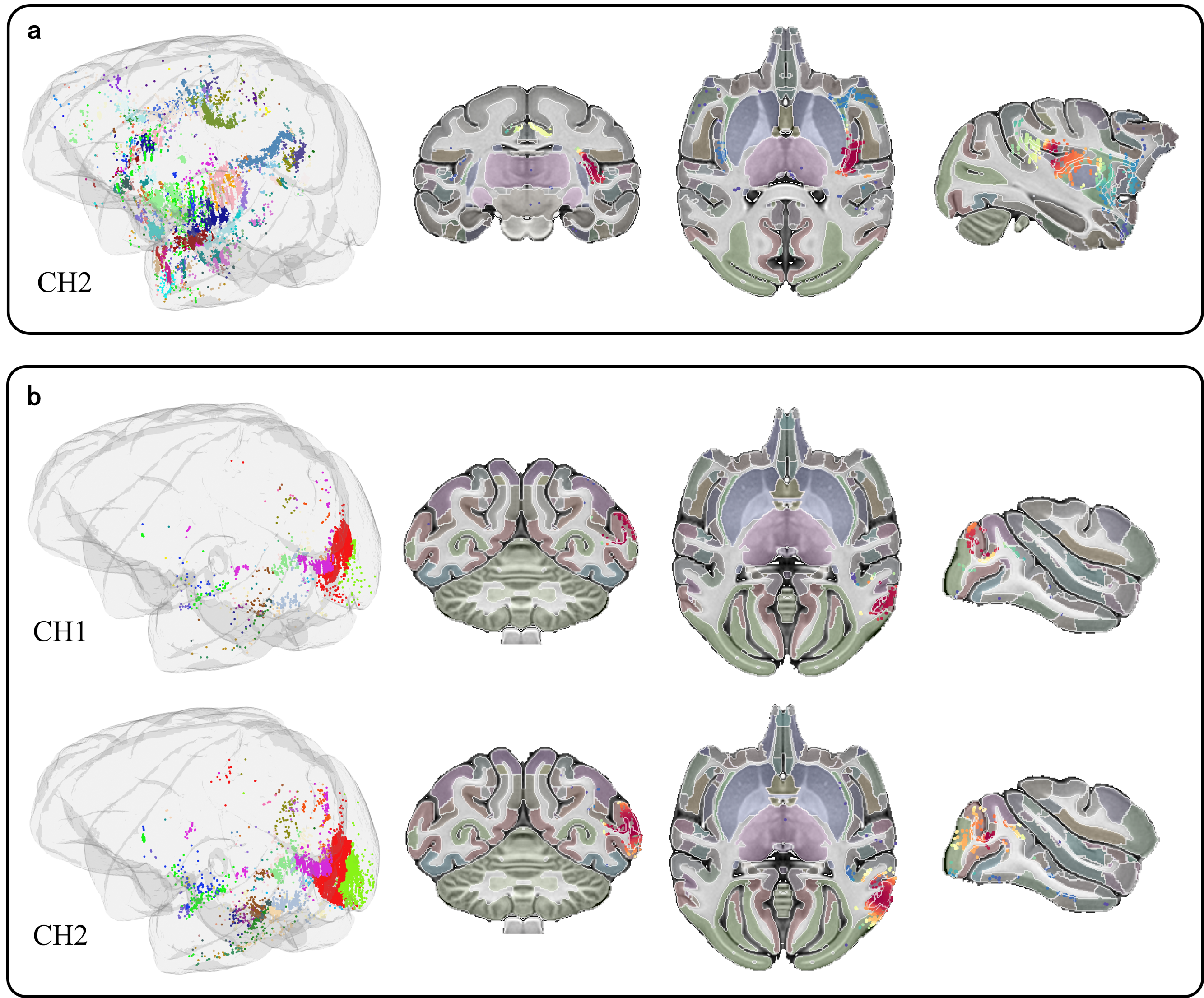

**Extended Data Fig. 7 | Distribution of soma in the standard space. a,** The injection site for channel 2 is in the id (insula) region. **b,** The injection sites for both channels are in the V1 primary visual region.

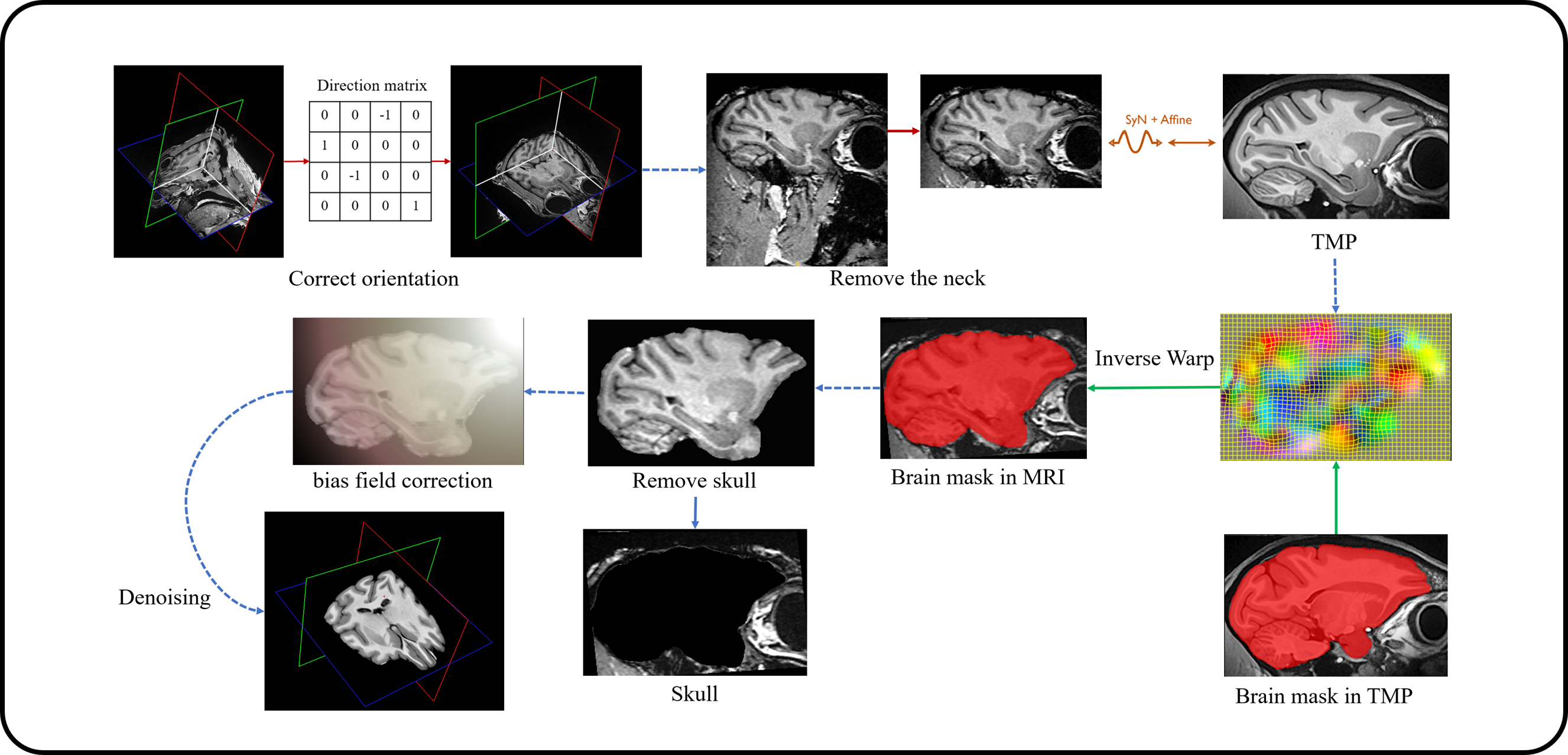

**Extended Data Fig. 8 | MRI preprocessing workflow.** The MRI preprocessing methods include brain mask extraction, bias field correction, and denoising, among others.

**Supplementary Table**

**Extended Data Table | Table of subregion merging relationships.**

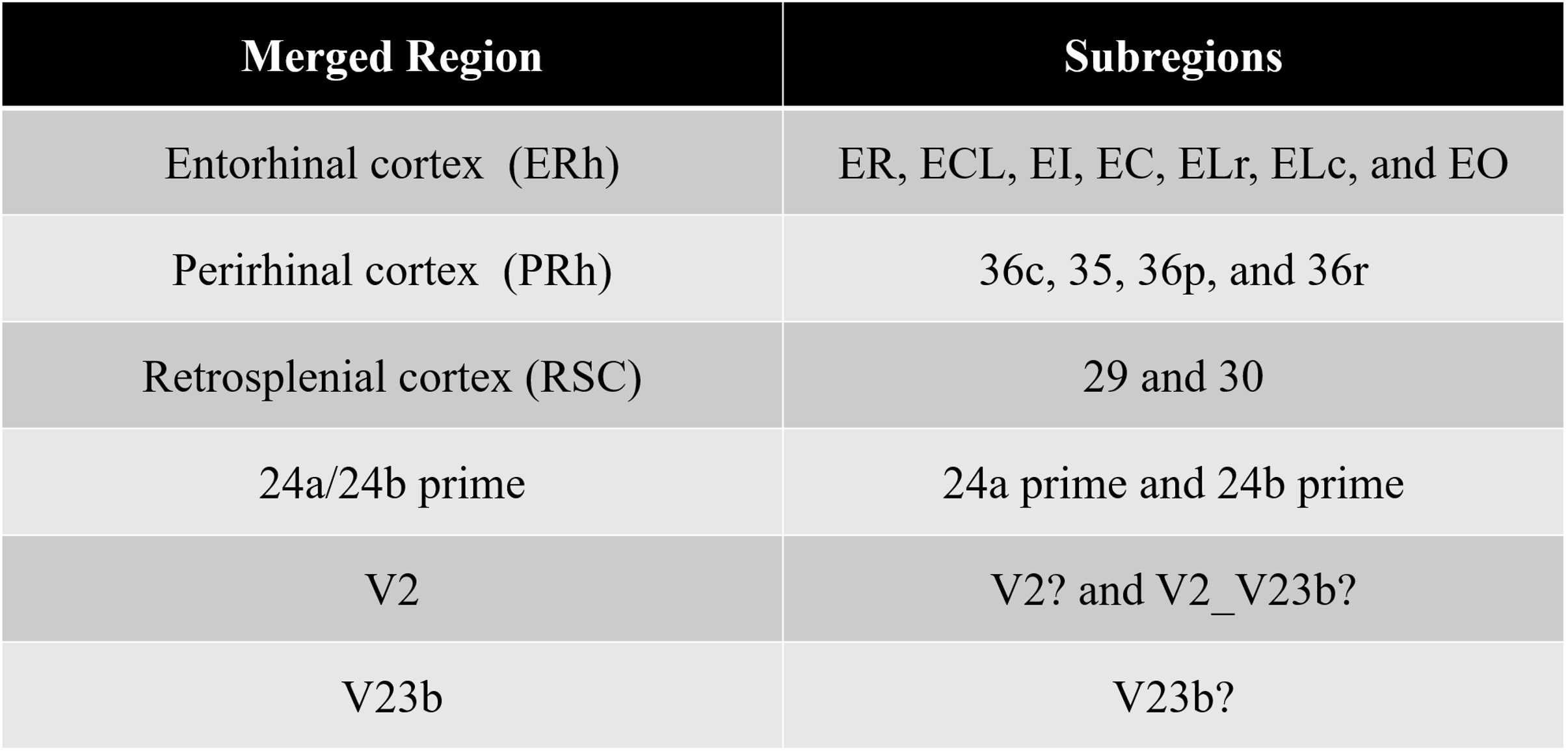
